# Aspirin Protective Effect Against Cyclophosphamide Hematological Toxicity In Experimental Animals

**DOI:** 10.1101/2021.06.29.450450

**Authors:** Imad Hashim, Zaid Al-Attar, Saba Jasim Hamdan

## Abstract

Bone marrow toxicity is the most important factor limiting the use of cytotoxic drugs like alkylating agents in cancer treatment. Recently PG synthase enzyme inhibitors have been reported to potentiate the cytotoxic effects of these agents on cancer cells but little is known if they can affect the toxicity of these agents on bone marrow or other tissues. Cyclophosphamide is one of the most commonly used alkylating agent.

In the present work, the effect of these PG synthase enzyme inhibitors, aspirin on cyclophosphamide myelotoxicity was determined employing the peripheral blood count to reflect bone marrow injury. The effect on body weight changes caused by cyclophosphamide was also determined.

1. Cyclophosphamide in doses of 25, 50 and 75 mg/kg i. v. produced as a dose dependent reduction in total WBC count, granulocyte, non granulocyte, and Hb% which was maximum on second day after injection and still present on 5th day post injection. It also produced a dose dependent reduction in body weight on day 5 after injection.
2. Aspirin in doges of 75, 150 and 300 mg/kg i. m. protected against the reduction in WBC counts ‘measured for 5 days after injection of cyclophosphamide (50 mg/kg). This protection was not dose dependent, though it was more optimum with 300 mg/kg and disappeared largely when a dose of 450 mg/kg was used. Aspirin did not prevent the changes in Hb% but retard the reduction in body weight caused by cyclophosphamide.
3. It is concluded that aspirin can help to reduce injury and enhance recovery from bone marrow toxicity caused by cytotoxic agents such as the alkylating drugs cyclophosphamide for which no specific antidote is available. Aspirin produces this effect possibly by eliminating the harmful inhibitory effect of excess PGs or leukotrienes, released by bone marrow injury on growth factors of haemopoietic progenitor cells.

The magnitude of this protection on WBC counts does not seem to differ between either PG synthase enzyme inhibitors or steroids when used alone or in combination although a synergistic effect in protecting erythropoiesis is observed.

## INTRODUCTION

Aspirin and related anti—inflammatory drugs inhibit metabolism of amino acids (AA) by cyclooxygenase only (1). Thus, they increased formation of leukotrienes by increasing the amount of AA that are available to lipoxygenase (2). Glucocorticoids inhibit phospholipase A3 activity so inhibit release of the potent, mediators PGs and leukotrienes. These mediators released by several stimuli contribute to genesis of sign and symptom of inflammation (3). Corticosteroids are more powerful as anti-inflammatory agents than aspirin-like drugs (4).

### A. WBC

Proliferation of normal granulocyte—monocyte progenitor is dependent on presence of colony-stimulating factors (5). PGE1 has inhibitory effect on both colony-stimulating factor and granulocyte-monocyte progenitor (6).

This inhibition of colony forming cell is dose-dependent and consistent with ability to increase intracellular level of CAMP (7).

PGE1 claimed to influence functions of B-lymphocytes selectively so depress humeral antibody response. PGs also affect T lymphocyte which are active in slowing tumor growth and killing malignant cells (8). PGE inhibit production and release of lymphokines by sensitized T-lymphocyte (9).

Leukotriene B4 (LT B4) is potent chemotactic for polymorph leucocyte, other leukotrienes do not share this property (10)

PGD2, 5-HPETE and 5-HETE enhance histamine release from human basophil (11) while PGE2 and PGI2 inhibit histamine release (12).

A predominant effect of PGs has been an inhibition of immunologic reactivity (13). Immunosuppressive activity of some tumors may be related to their ability to produce PGs (14).

### B. Platelet

PGE1 and PGD2 are inhibitors of aggregation of human platelet. PGI2 is 50 times more potent inhibitor.

PGE2 is stimulator at low concentration and inhibitor at high concentration.

TX A2 is powerful inducer of platelet aggregation and platelet release reaction. No evidence available for direct involvement of PGs in formation of platelets from megakaryocytes (15).

Effect of PGs on platelet aggregation have been implicated in hematogenous metastasis of tumors.

Infusion of PGI2 before or after injection of tumor cell markedly inhibit establishment of tumor colonies; this is attributed to inhibition of platelet aggregation rather than vasodilation (16).

### C. RBC

PGE1,E2 at low concentration decrease fragility of RBC, at high concentration increase fragility (17).

PGE2 may interfere with red cell maturation and may reduce amount of synthesized Hb that it contain (18).

### CYCLOPHOSPHAMIDE

Among the alkylating agents, the most widely used in therapy of both experimental and human cancer is cyclophosphamide.

Cytotoxic agents have produced beneficial results in cancer chemotherapy, particularly on hematological and lymphoid malignancies. However, the toxic effects on normal body tissues, especially bone marrow, is the main limiting factor in their use. Many procedures have been used to reduce toxicity of cytotoxic agents on normal body tissues in order to permit the use of higher doses safely, while also increasing the rate of tumor cell kill. Among the agents used to modify cyclophosphamide toxicity are N—acetyl cysteine (19) and mesna (20).

In our study, we employed cyclophosphamide since it is a widely used alkylating agent in cancer chemotherapy and immunosuppression.

### AIM OF STUDY

Several doses of aspirin will be employed to investigate their effect modifying cyclophosphamide toxicity on several hematological parameters in addition to total body weight.

## MATERIALS AND METHODS

### ANIMALS

Adult male rabbits, weighing 1-2 kg., were allowed free access to water and commercial food pellets through all days of the experiment.

### DRUGS

Drugs were freshly prepared for each experiment. The following drugs were employed

1. Cyclophosphamide (Endoxan-Baxter;Baxter Oncology GmbH Kantstrasse2 D-33790 Halle, Germany, Vials of 200 mg/10cc)
2. Aspirin (Aspegic: from the laboratories synthelabo, France; Vials of 500 mg/ 5 cc).
3. Heparin (Heparin Leo: from Leo pharmaceutical products, Ballerup— Denmark, Vials of 25000 unit/ 5 cc). It was given to rabbits in doses of 5000 unit/ kg i. v. prior to taking blood samples.

### Hematological Indices Employed

Blood samples were taken from animals and analyzed to get complete blood count (in terms of RBSs, WBCs, platelets, and hemoglobin) using automated electronic counter (Hematology auto-analyzer - BECKMAN COULTER, ACT. 5 diff. USA).

#### PROCEDURE

##### 1. INTERACTION OP DIFFERENT DOSES OF ASPIRIN WITH CYCLOPHOSPHAMIDE (50 mg/kg)

Four groups, each has 6 rabbits included, that received aspirin i.m. in doses of 75, 150, 300 and 450 mg/kg, respectively, daily for 3 days. The second injection of aspirin in those groups was given simultaneously with cyclophosphamide 50 mg/kg i.v. Blood samples were taken from the marginal ear vein before the first aspirin injection and daily for five days after cyclophosphamide injection, for the estimation of Hb%, WBC counts (total and differential) and platelet.

Since the absolute WBC or platelet counts vary remarkably between animals, percentages were included to avoid this variation between animals by considering the pre-treatment value of each animal as 100%. The count of total granulocytes was taken since heparin clearly interferes in differentiating clearly between them, i.e. neutrophils, eosinophils or basophils. However, since neutrophils form the largest proportion of granulocyte, alterations in this count probably largely reflects changes in neutrophil count. Similarly, heparin interferes in clearly differentiating between non granulocytes (lymphocytes and monocytes). However, since lymphocytes comprise the majority, changes in total non-granulocytes count probably largely reflects variations in lymphocyte counts.

#### STATISTICS

Statistical comparisons between treated groups and their respective controls was done with ANOVA test. Significance was taken to reject the null hypothesis when P<0.05. To determine the dose-dependent effect of drugs, the Dunnett’s multiple comparisons test was conducted.

## Results

### EFFECT OF DIFFERENT DOSES OF ASPIRIN IN CYCLOPHOSPHAMIDE (50MG/KG) TREATED RABBITS

The parameters that we investigated have shown a great variation throughout the period of study i.e. (the 5 days of subjecting the lab animals to the effect of cyclophosphamide). Thus, we found that the most practical and the most conclusive approach is to depend on the results of the 5^th^ day (the end of the study). Another important point is that significance level that is mentioned in results is regarding one direction. i.e. the positive change regarding the studied parameters. E.g. in **figure 2.A** we found that aspirin in doses of 75mg/kg significantly reduced WBC count (P <0.01) compared to control. Nevertheless, this significant effect is in negative direction i.e. WBC count decreases which is an undesirable effect.

**Figure.1:**
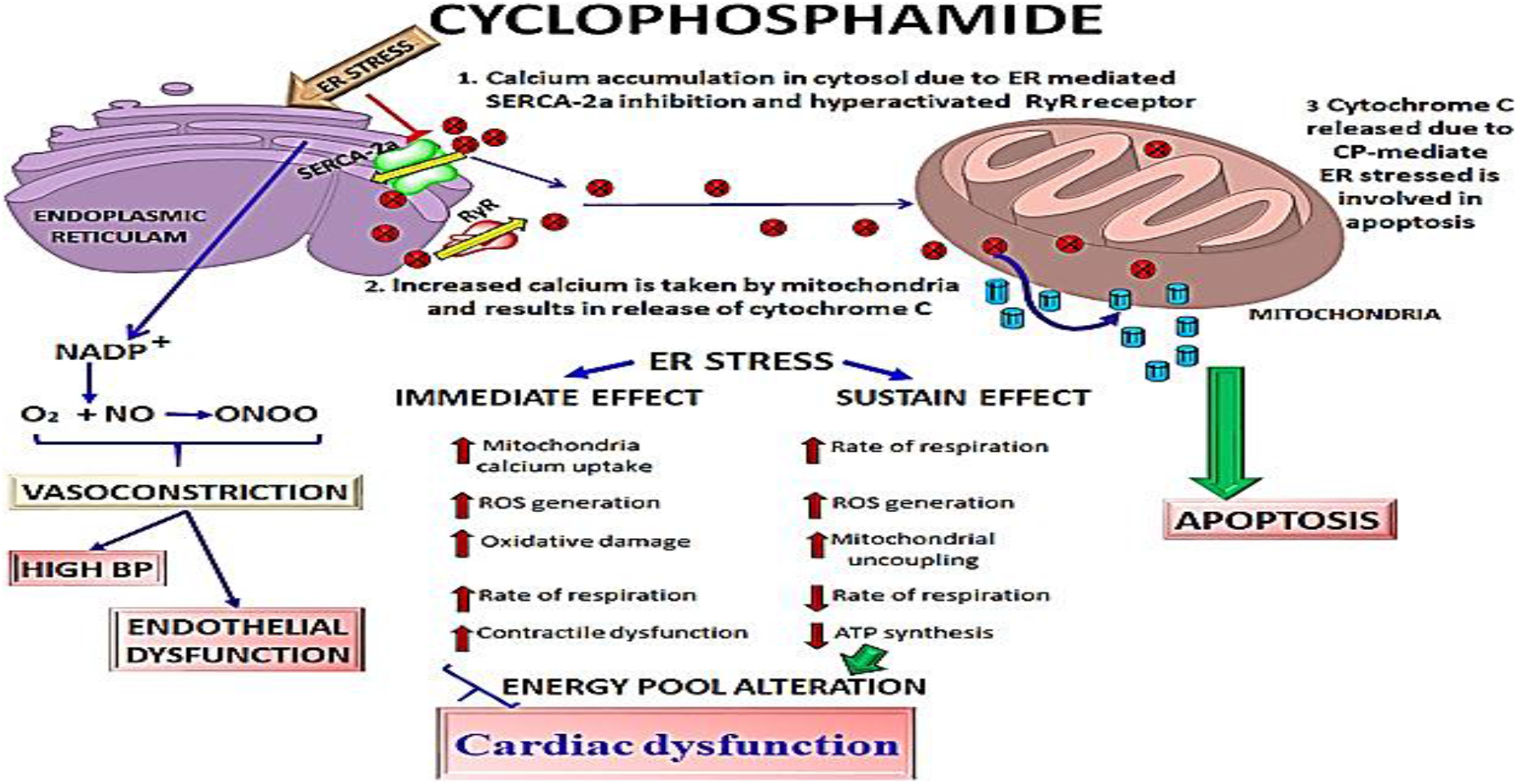
The role of cyclophosphamide in the treatment of malignancy.

**Figure (2).**
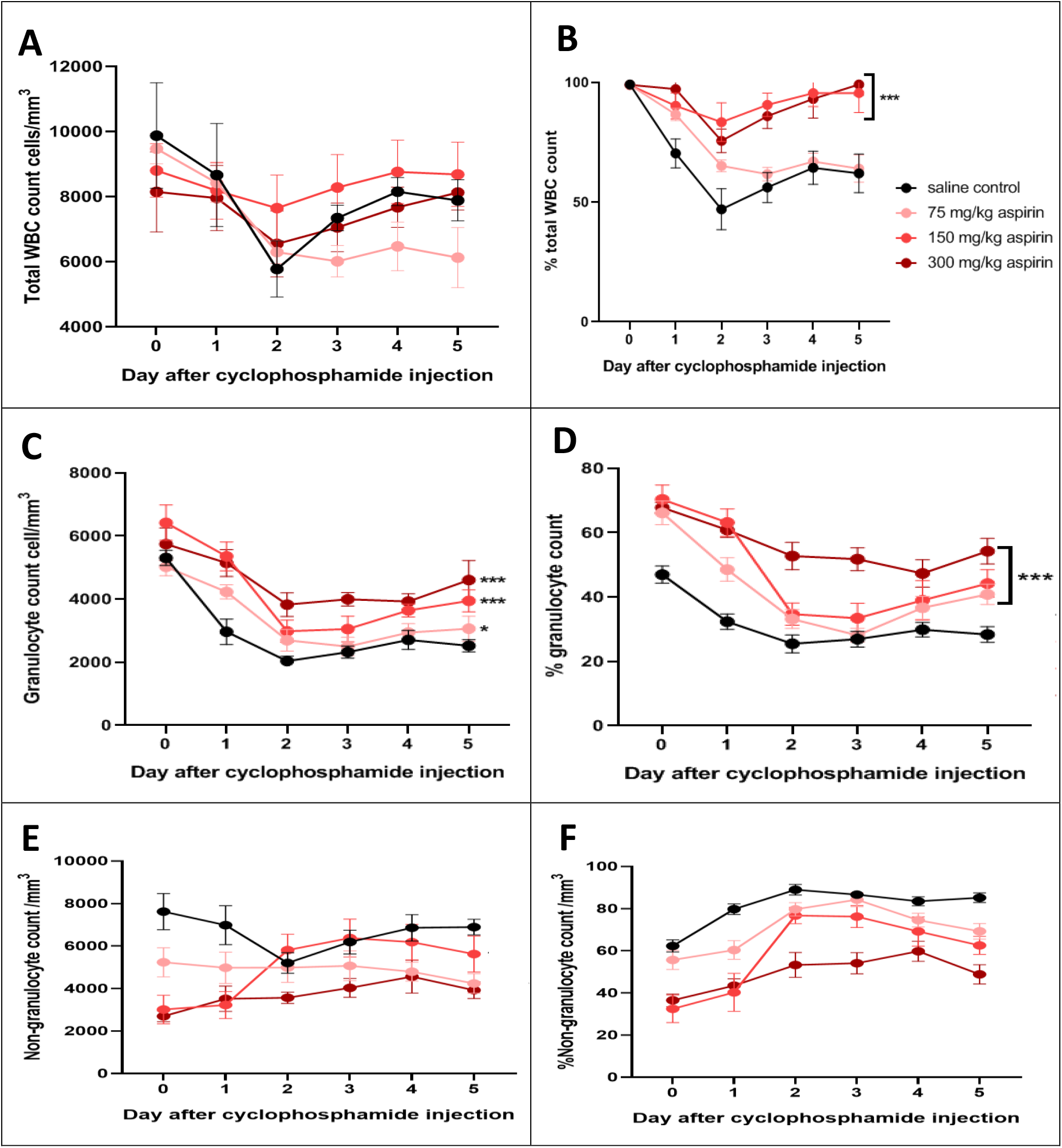
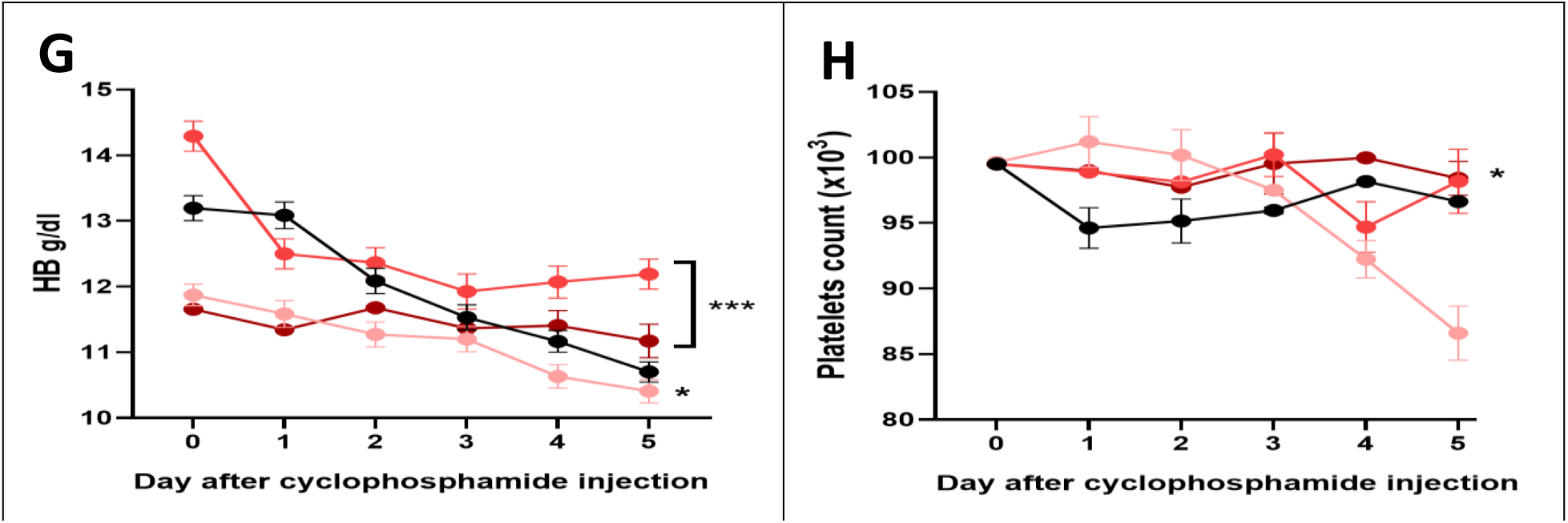
Effect of different doses of aspirin (75, 150 and 300 mg/kg i .m.) on blood picture in cyclophosphamide (50 mg/kg I.V.) treated rabbits. Effect On total WBC count. A: absolute, B: percentage. Effect on granulocyte count. C: absolute, D: percentage. Effect on non— granulocyte count. E: absolute, F: percentage). Effect on G: Hb%, H: platelet count percentage. *:p<0.05, **:p<0.01 and p:<0.001 according to ANOVA test and Dunnett’s multiple comparisons test.

Thus, we looked for changes in the positive direction for 150 and 300mg/kg. Findings in terms of these two doses are not significant. However, they reflect an ameliorating effect for the toxic effects of cyclophosphamide.

Aspirin improved the relative count of WBC in cyclophosphamide treated lab animals at all doses of aspirin with high statistical significance for the 150 and 300mg/kg as shown in **Figure 2.B**.

Aspirin at has shown to be protective against cyclophosphamide induced granulocyte lowering effect (absolute count and percentage) at all doses with high statistical significance especially for 300mg/kg dose (**figures 2C** and **2D)**.

However, Aspirin failed to improve the absolute and relative non granulocyte counts as shown in **figures 2.E** and **2.F**.

Aspirin improved hemoglobin levels at 150 and 300 mg/kg doses with high statistical significance while the 75mg/kg dose led to negative effect on hemoglobin level as shown in **figure 2.G**.

Platelet count improved significantly for 300 mg/kg dose and non significantly for 150mg/kg dose. While the 75mg/kg dose had a negative impact on platelet count as shown in **figure 2.H**.

## Discussion

The hematopoietic system is very susceptible to effects of nitrogen mustards including cyclophosphamide (21). Bone marrow depression is the most important limiting factor for using these agents in cancer treatment. Being a chemical injury to the bone marrow, cyclophosphamide is associated with de novo synthesis and release of various PGs, which could have a further adverse effect on hemopoiesis colony stimulating factors synthesized in situ by T-lymphocytes and macrophages which is also essential for proliferation and maturation of granulocytes and non granulocytes (22).

### CSFs include

Granulocyte colony stimulating factor (G—CSF), Granulocyte— macrophage colony stimulating factor (GM—CSF), Macrophage colony stimulating factor (M—CSP) and Interleukin—3 (IL 3).

G-CSF and GM-CSF cause major, short term, dose—dependent increase in neutrophil counts after intravenous infusion, in addition GM-CSF increase absolute counts of monocytes, eosinophils and lymphocytes (23).

CSFs will almost certainly be used in combination, for the amelioration of bone marrow toxicity and mucositis associated with intensive chemotherapy (24).

It is thought that concurrent employment of these factors will allow the use of large doses of cytotoxic agents in cancer treatment to obtain larger cells kill with minimum toxicity and also might help to prevent the emergence of drug resistance among susceptible cells.

Alternatively, the use of agents which would increase the endogenous level of these growth factors by enhancing their production and/or reducing their catabolism, or potentiating them by removing any endogenous inhibition upon their action might achieve the same results.

Since PGs are known to be inhibitory to these growth factors, the use of PGS inhibitors such as aspirin might be among possible ways of achieving this aim in protecting against bone marrow injury and allowing greater selectivity in use of cytotoxic agents.

In our work, we employed cyclophosphamide since it is one of the most widely used cytotoxic alkylating agents. In addition, its bone suppressive effect might apply to bone marrow injury by other cytotoxic agents whether chemical or physical (e.g. irradiation).

In our work, different doses of cyclophosphamide caused significant depression in total WBC count in a dose-dependent manner which was maximum on the 2nd day after injection. Recovery of haemorgam which was evident on the 5th day after injection probably reflects the disappearance of the drug which has a half—life of 4-6 hrs, and resumed activity of resistant stem cells and progenitor cells.

This reduction is mainly due to a significant decrease in granulocytes more than non-granulocytes. Since the predominant granulocytes are the neutrophil, this probably represents a more prominent damage to neutrophil precursors in bone marrow by cyclophosphamide metabolites producing a dose-dependent alkylation of myeloblast and myelocyte.

In terms of total WBC count, aspirin has shown an ameliorating effect at 150 and 300 mg/kg doses, however, it is not significant. While, in terms of WBC percentage it there is a significant ameliorating effect at these doses and being maximal at 300 mg/kg.

PGE2 was found to be a dose-dependent inhibitor of proliferation of normal colony-forming cells (25) and this is consistent with its ability to increase intracellular level of CAMP (26) which would produce considerable lowering of granulocyte precursor cell.

In our work, reduction of PGs synthesis and release by the PGs inhibitor; aspirin might reduce their harmful effect on haemopoietic precursor cells in bone marrow. This explains the protective role against reduction of granulocyte, particularly neutrophils, mediated by cyclophosphamide.

On the contrary, aspirin has shown an exacerbating effect on non-granulocyte counts and percentage. The differential counts of granulocytes and non-granulocytes are showing mirror-image changes that reflect and confirm the validity of our results.

Cyclophosphamide also produced a dose-dependent decrease in Hb level. The red cell precursors may be affected to the same degree by alkylation as the precursors of granulopoiesis in bone marrow (27).

It has been found that there is a direct relationship between renal prostaglandin (PGE2) and erythropoietin levels. Inhibition of prostaglandins synthesis by aspirin effect on cyclo-oxygenase may lower erythropoietin levels (28). Nevertheless, inhibition of cyclo-oxygenase would shift the balance toward leukotrienes production. The leukotrienes (LT) B4 and C4 caused a reduction in granulocyte-macrophage as well as erythroid colony numbers in a dose-dependent manner (29). Thus, it seems that the effect of leukotrienes is more powerful than prostaglandins in our study conditions.

As most cytotoxics, cyclophosphamide is well known for causing thrombocytopenia. However, the use of aspirin has shown a dramatic significant protective effect for platelets as shown for 300mg/kg dose.

It seems that a similar mechanism is also protecting the non-granulocyte, i.e. lymphocyte and monocyte, and our PGSIs are probably mainly acting by eliminating these harmful effects of PGs on leucopoiesis.

However, the action of aspirin in inhibiting PG synthesis does not seem to be enough to protect against alkylation and injury to erythroid precursor in bone marrow by cyclophosphamide metabolites, with subsequent lowering of hematocrit or Hb%. Although post block they almost completely stops PG production by cyclooxygenase, their use is associated with greater availability of AA for lipoxygenase with subsequently greater production of leukotrienes (2). It is possible that high level of leukotrienes can affect bone marrow erythropoiesis as adversely as PGs.

Other possible mechanisms that might be possibly involved in the protective effects of PGSIs on leukopoiesis and in reducing cyclophosphamide toxicity include aspirin protein binding. Aspirin is extensively bound to plasma proteins (30). Cyclophosphamide is bound to plasm proteins to less extent (31) and thus being both administered would increase the unbound fraction of cyclophosphamide. Possibly a similar competition occurs at the cell level thus reducing entry of cyclophosphamide into cells.

In conclusion, it seems that aspirin can help to reduce injury and enhance recovery from bone toxicity caused by cytotoxic agents such as the alkylating drug cyclophosphamide for which no specific antidote is available. Aspirin produces this effect possibly by eliminating the harmful inhibitory effect of excess PGs or leukotrienes (released by bone marrow injury) on growth factors of haemopoietic progenitor cells. It is speculated that treatment with these agents in future would increase the selectivity and safety of cytotoxic agents, allowing larger doses to be used to increase cancer—cell kill and probably prevents emergence of drug resistance.

